# Sensory Cue Combination in Children Under 10 Years of Age

**DOI:** 10.1101/501585

**Authors:** James Negen, Brittney Chere, Laura Bird, Ellen Taylor, Hannah E. Roome, Samantha Keenaghan, Lore Thaler, Marko Nardini

**Author notes:** Declarations of interest: none.

## Abstract

Cue combination occurs when two independent noisy perceptual estimates are merged together as a weighted average, creating a unified estimate that is more precise than either single estimate alone. Surprisingly, this effect has not been demonstrated compellingly in children under the age of 10 years, in contrast with the array of other multisensory skills that children show even in infancy. Instead, across a wide variety of studies, precision with both cues is no better than the best single cue – and sometimes worse. Here we provide the first consistent evidence of cue combination in children from 7–10 years old. Across three experiments, participants showed evidence of a bimodal precision advantage (Experiments 1a and 1b) and the majority were best-fit by a combining model (Experiment 2). The task was to localize a target horizontally with a binaural audio cue and a noisy visual cue in immersive virtual reality. Feedback was given as well, which could both (a) help participants judge how reliable each cue is and (b) help correct between-cue biases that might prevent cue combination. A feedback effect was found in Experiment 2, with children who were given single-cue feedback showing the highest rate of cue combination. Given this, we suggest that children at 7–10 years old are capable of cue combination in principle, but must have sufficient representations of reliabilities and biases in their own perceptual estimates as relevant to the task, which can be facilitated through task-specific feedback.

## 1. Introduction

Cue combination is when two independent noisy perceptual estimates are merged together as a weighted average, creating a unified estimate that is more precise than either single estimate alone (Ernst & Banks, 2002). This process is a key aspect of Bayesian Decision Theory (Maloney & Mamassian, 2009), allowing people to deal in a near-optimal way with the different sources of sensory noise involved in perceptual decisions. For example, when both looking at and holding an object, we have information about its size from vision as well as from touch. If we always take the approach of averaging the available estimates while weighting each independent estimate by its precision, we will always arrive at the most precise possible estimate of size. The main alternatives would be strategies such as cue selection, where one cue is used and the other is ignored, or more rote forms of integration like taking an unweighted average of the estimates.

Since a seminal report in 2002 (Ernst & Banks, 2002), evidence has accumulated that a Bayesian cue combination model is a good description of how adults typically perform when given multiple independent cues to a singular perceptual judgement (Knill & Pouget, 2004; Pouget, Beck, Ma, & Latham, 2013). In particular, the strongest single test for cue combination is a bimodal precision advantage. If two cues are being combined (averaged) in a Bayes-like manner when they are both available, judgements should be more precise with both cues available than the best single cue alone. However, nearly all developmental studies so far suggest that children up to the age of 10 years do not show this effect (Adams, 2016; Burr & Gori, 2012; Dekker et al., 2015; Gori, Del Viva, Sandini, & Burr, 2008; Jovanovic & Drewing, 2014; Nardini, Bedford, & Mareschal, 2010; Nardini, Begus, & Mareschal, 2013; Nardini, Jones, Bedford, & Braddick, 2008; Petrini, Remark, Smith, & Nardini, 2014). This is despite using a broad range of tasks – uni-modal (different cues to depth but all within the visual modality) as well as multi-modal (e.g. vision and touch); two-alternative forced-choice as well as more naturalistic responses (e.g. point where it is). While this provides compelling evidence of late development, the mechanisms underlying this late development remain unclear.

Here we answer a simple but critical question about the development of cue combination, shaping the direction of future theory. Should we think of the emergence of cue combination as a relatively unified and sudden event at about 10 years old? If so, we might favour unified explanations such as biological changes at the beginning of puberty driving its emergence. Alternatively, does cue combination emerge at meaningfully different time points for different tasks? If so, we should develop a nuanced theory that pins the emergence of different aspects of cue combination to dissociable causes contained within the earlier-emerging tasks. Clarity on this point is a key step towards a full understanding of how cue combination develops and what drives its development.

The potential applications of this knowledge range from the treatment of sensory issues in childhood over to robotics and sensory augmentation. The human brain and perceptual systems are self-organizing entities that adapt in complex ways to the needs of the environment throughout development (Stoianov & Zorzi, 2012; Van Orden, Holden, & Turvey, 2003). Understanding how the system organizes itself to implement sophisticated new algorithms – e.g. those approximating Bayesian inference (Pouget et al., 2013) – provides the necessary theoretical backdrop to understand sensory issues in atypical populations such as autism and schizophrenia, in which these computations may operate differently (Montague, Dolan, Friston, & Dayan, 2012; Wang & Krystal, 2014). Further, creating human-like adaptive capabilities in robots also requires a thorough understanding of the course of human development (Lake, Ullman, Tenenbaum, & Gershman, 2017). Finally, knowing how children learn to integrate typical cues will help us understand how to help adults integrate augmented sensory devices into their perceptual systems – a feat which adults don’t perform just by exposure to a new device’s use (Goeke, Planera, Finger, & König, 2016). To move towards answers that will help with these applications, we tested two hypotheses about the course of emerging cue combination skills in development, which make opposing predictions.

### 1.1. The Hard Limit at 10–11 Hypothesis

This hypothesis states that children under 10 years old have a fundamental block, a relatively ‘hard’ limit, which prevents them from combining cues across all of the different tasks given. This leads to a relatively unified emergence of cue combination at 10–11 years old. This is a relatively parsimonious explanation that still fits the vast majority of the available data. To anchor this discussion, it is necessary to look in some detail at a recent study of how visual depth cues, specifically binocular disparity and relative motion, develop in middle/late childhood (Dekker et al., 2015). This study stands out as having a high number of participants (N=142), sampled continuously from a relatively narrow age range (6–12 years), and with each participant completing an unusually high number of trials for a developmental sample (360). Three measures of cue combination were obtained (two behavioural and one from fMRI). All three were treated with a regression, finding a divergence from no effect after the 10^th^ birthday but before the 11^th^. All three showed no significant cue combination effect in children under 10.5 years old, but a very clear and significant effect in children over 10.5 years old. This provides convergent, consistent evidence that the combination of visual depth cues is present after 11 years but not present before 10 years, at least under their task parameters.

This finding of a transition at about 10 years also converges with other findings from studies that looked at the combination of different cues. There is no finding of a bimodal precision advantage under 10 years in a wide variety of studies with different modalities and different types of judgements: self-motion and landmark cues to navigation (Nardini et al., 2008), visual and haptic cues to size and orientation (Gori et al., 2008), disparity and texture cues to slant (Nardini et al., 2010), visual and auditory cues to spatial and temporal bisection points (Burr & Gori, 2012), visual and haptic cues to object size (Jovanovic & Drewing, 2014), and finally visual and auditory cues to the number of events (Adams, 2016). Some of these studies did not test children shortly after the age of 10 years, but those that did found a transition shortly after. For example, a partial integration model fit the majority of participants at 10–11 years (Adams, 2016) and a bimodal precision advantage was seen at 12 but not 10 years (Nardini et al., 2010).

The idea of a unified transition at about 10 years also fits with a proposed theory of why cue combination would not be seen in young children (Burr & Gori, 2012). It might actually present a subtle but important advantage. This theory suggests that failing to combine cues might make it easier to use cues for cross-calibration. In this context, cross-calibration is defined as the elimination of relative biases. A relative bias is when a given state of the world, cued in two different ways at different times, leads to different average judgements of the world state (e.g. a light and sound straight ahead are judged on average as 5 degrees left and 5 degrees right, respectively, which is a relative bias of 10 degrees). This theory also states that cross-calibration needs are most intense during the first ten years of life when the eyes, ears, and skull are developing the fastest. If these premises are true, then combining cues from an early age might bar children from effectively eliminating relative biases, leading to an extended period of disjointed multimodal perception. In that sense, it could be advantageous for children under 10 years to keep cues separate rather than combine them.

Another reason to favour a unified transition account is the possibility that the biological substrates implementing reliability-weighted averaging are not capable of accurately carrying out these computations until near the start of puberty. These mechanisms are not yet well understood, but one account proposes that they depend on the balance of excitation and inhibition over relatively large populations of neurons (Ohshiro, Angelaki, & DeAngelis, 2011). Their development is even less well understood, but it is certainly possible that it is protracted during the childhood years and into adolescence, like many aspects of cortical organisation and function (Gogtay et al., 2004).

A recent study with adults also fits with this type of explanation (Negen, Wen, Thaler, & Nardini, 2018). Adults were asked to learn new depth cues – based either on sound delay or wrist vibration – and combine these with a noisy visual cue. This means that participants had a cue-specific experience that was child-like (i.e. low), but still enjoyed the full neural maturation of adulthood. If the level of cue specific experience was more important, we would expect child-like performance from them. Instead, participants learned to combine either new cue with the noisy visual cue in just a few hours, emphasizing a strong role for general neural maturation in predicting cue combination. This is consistent with the idea that children might simply need to achieve the neural changes involved in the first decade of life before they can combine cues.

### 1.2. The Undiscovered Task Hypothesis

The alternative hypothesis is that children who are under 10 years old actually can combine cues on the right task with the right parameters, but this task has not yet been discovered yet. Under this view, children at 7–10 years old are fundamentally capable of combining cues if circumstances allow and they are supported towards that. This would mean that development is less like a hard limit at 10 years old and more like a gradient of earlier- and later-emerging cue combination performance on different tasks. This hypothesis has not yet seen any reliable support in the existing literature – by our count, only one (Nardini et al., 2013) out of 26 tests for a bimodal precision advantage below 10 years has been positive, a proportion not greater than the expected Type I error rate. However, it remains the obvious alternative to existing theory (Burr & Gori, 2012; Ernst, Rohde, & van Dam, 2016). Our central approach is to see if a new task can provide consistent evidence of cue combination in this age range.

### 1.3. The Present Study

The key evidence required to favour the Undiscovered Task Hypothesis over the Hard Limit Hypothesis is a robust, repeatable finding that children under 10 years old can combine cues, standing in contrast to previous studies having shown combination only at 10+ years. To achieve this, we chose a new task with different parameters than those used in previous studies. Specifically, we chose to focus on the parameter of feedback as way of making a new task, in essence investigating how variations in feedback affect cue combination behaviour. Feedback might help participants understand the relative reliability of their perception of the two cues, which is the key to setting the correct weights (Ernst et al., 2016), and poor weight choices in this age range have been empirically demonstrated several times (Gori, Sandini, & Burr, 2012; Nardini et al., 2010). Feedback might also help reduce any relative biases. In adults, a strong relative bias prevents cue combination, since it leads to the perception of two separate causes (Shams & Beierholm, 2010). For example, it does not make sense to average the perceived positions of a light and a sound source, creating a unified estimate of their single cause, if a strong relative bias leads to the inference that the cues are from different places. The Hard Limit Hypothesis states that this should all be irrelevant and we still should not see a bimodal precision advantage in children younger than 10 years. The Undiscovered Task Hypothesis allows for the possibility that this effect will be found in this age range.

For Experiment 1a, we ran a variant of a previous spatial audio-visual cue combination task with 7–10 year old children (Gori et al., 2012). The previous study found a visual capture effect (only the visual information was used) in their version without feedback. We altered Gori et al.’s paradigm by giving feedback on single-cue trials (audio-only or visual-only), but not the audio-visual trials, forcing participants to infer how to combine the two cues only from their experience with each cue in isolation (Maloney & Mamassian, 2009). To make the feedback more meaningful, we also moved from a two-alternative forced-choice (2AFC) bisection paradigm to an absolute judgement paradigm; feedback on absolute judgements should, in principle, be more informative than feedback on 2AFC judgements. In brief, our task asked participants to judge the location of a target on a left-right horizontal axis in front of them, relying either on a noisy visual cue, a spatialized sound, or both. The target was ‘hiding’ behind an opaque wall and they used a computer mouse to click where they thought it was on the horizontal axis. The key analysis is whether judgements are more precise with both cues available than the same participant’s best single cue. This is a result only expected under cue combination (weighted averaging).

The results of Experiment 1a were very surprising, showing a bimodal precision advantage. Given the large number of studies on this (reviewed above), almost all of which found null results in this age range, we wanted to make sure the results were replicable (i.e. not simply a Type I error due to a large number of independent tests across many labs and studies). We therefore ran Experiment 1b, which was a direct replication of Experiment 1a, finding the same pattern of results.

For Experiment 2, we wanted to look at this phenomenon with a slightly different approach to see if we could find converging evidence. We reasoned that if children at this age are combining cues in a task like this, we should not only see a robust bimodal precision advantage, but a model comparison analysis should also generally favour a cue combination model over any single-cue models as well. Experiment 2 also had two secondary aims. The first was to see if there is an effect of feedback that cannot be explained by simple task engagement. To achieve this, we did not compare full-feedback and no-feedback conditions, as any failures to combine cues in the latter could always be because they simply find the task less engaging. Instead, we varied the type and amount of feedback across conditions in a manner that did give some feedback to all participants in order to keep motivation and engagement high. The second was to see if children can adapt to varying levels of visual reliability on a trial to trial basis. This is expected under the full remit of Bayesian Decision Theory (Maloney & Mamassian, 2009), but not under current models of multisensory learning in development (Daee, Mirian, Ahmadabadi, Brenner, & Tenenbaum, 2014; Weisswange, Rothkopf, Rodemann, & Triesch, 2011). Throughout experiments 1a, 1b, and 2, our primary motivation was to see if children at 7–10 years combine cues in our new task, which would discredit the Hard Limit Hypothesis in favour of the Undiscovered Task Hypothesis.

## 2. Experiments 1a and 1b

In Experiment 1a, 7- to 10-year-old children were tested on their visual and/or auditory localisation in the horizontal plane in an immersive virtual reality (VR) environment. Their task was to find a tiny virtual woman in a hot air balloon (‘Piccolina’) hiding behind a wall with the aid of a brief visual cue, a spatialized sound, or both (Figure 1). To show where they perceived Piccolina to be, children used a virtual tool that they controlled using a computer mouse. This allows us to analyse the data with standard statistical tests to compare unimodal precision and bimodal precision (Ernst et al., 2016). Based on the Hard Limit hypothesis, we would predict that their precision with both cues available would be no better than the same child’s precision with their best single cue. Based on the Undiscovered Task hypothesis, we would predict that precision with both cues together would be higher on average than with the best single cue. Since the results of Experiment 1a stood out in contrast to many previous similar studies, Experiment 1b directly replicated Experiment 1a.

### 2.1. Method

#### 2.1.1. Participants

77 participants were initially tested in total across Experiments 1a and 1b. Nine cases were excluded, as participants failed to make estimates above chance in at least one condition. This left 68 participants for analysis (33 males, 35 females). Ages ranged from 7-years 3-months to 10-years 9-months, with nine 7-year-olds, twenty 8-year-olds, twenty-six 9-year-olds and thirteen 10-year-olds. Two phases of data collection occurred: 29 participants in Experiment 1a, and 39 participants in Experiment 1b. All children were enrolled in Primary Schools in County Durham, UK. All children had normal or corrected-to-normal vision and hearing. To the knowledge of the researchers, no children had been diagnosed with any perceptual or developmental disorder that might have affected task performance. This study was approved by the Psychology Department Ethics Sub-Committee at Durham University (reference: PGT2018MN1). Parents gave written informed consent.

#### 2.1.2. Apparatus

Virtual Reality technology was used. Visual motion and positional cues in virtual space are processed similarly to how they would be in real space (Foreman, 2009), allowing for ecological validity whilst retaining a high level of experimental control. This study used WorldViz Vizard 5 software and the Oculus Rift headset. AKG K 271 MK II headphones were used with a SoundBlaster SB1240 sound card.

A simple virtual environment was created with four notable features (Figure 1). There was a large curved wall 2m away from where the participant sat. It had a gray fixation circle at its center. A red dot, when shown, was under the participant’s control and used to indicate their estimates. The target character (‘Piccolina’) was a 5 cm tall woman in a 15 cm tall multi-colored hot air balloon that could either jump excitedly and clap her hands or lean over the side as if in tense anticipation of the coming events. Participants were essentially trying to estimate where Piccolina was hiding behind the top edge of the wall.

**Figure 1.**
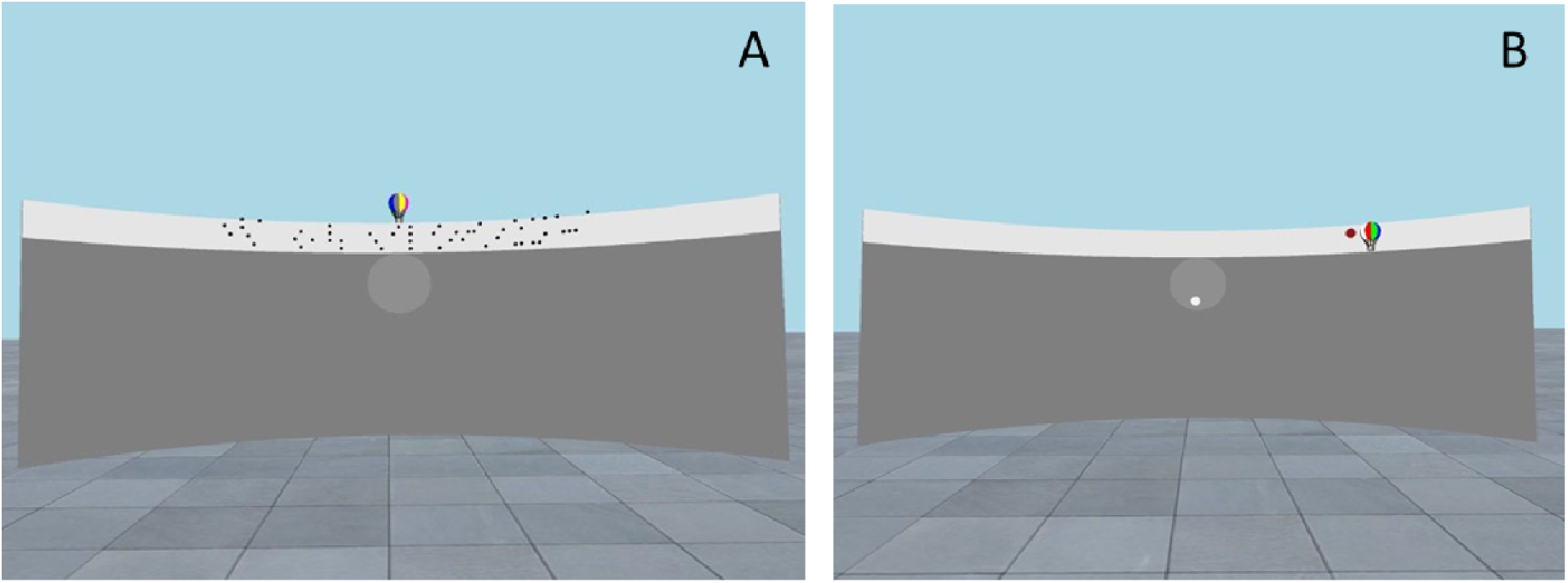
Screenshots from the task. Panel A is from the Demonstration phase (described fully in *Procedure*), where participants were familiarized with the visual stimulus. Panel B shows example feedback from the Data Collection phase where the participant has placed the response (red dot) about 75% of the way to the right and the correct target (‘Piccolina’ in the hot air balloon) was very near, slightly further right.

#### 2.1.3. Stimuli

Visual stimuli consisted of an array of 64 spheres with a radius of .008m, placed uniformly along the horizontal axis of the wall and randomly along the vertical axis (Figure 1A). During 250ms, spheres moved 1/7^th^ of the way to the target location before disappearing. This visual cue to location was designed to be uncertain, giving a potential advantage for participants to combine the visual cue with the auditory cue when both were available. Judging the point of convergence can be described as a motion integration or coherent motion task (Burr & Thompson, 2011).

The audio stimuli consisted of white noise amplitude modulated at 10Hz. This lasted for 300ms. Audio stimuli were spatialized using the ‘small pinnae’ head-related transfer function from the CIPIC head-related transfer function (HRTF) database (Algazi, Duda, Thompson, & Avendano, 2001) for 117 different points along the wall. For these experiments, audio and visual stimuli were both accurate and aligned (i.e. no perturbations were shown). All variation in results thus reflected internal noise and biases.

Audio-spatial stimuli typically show measurable biases, even when using a speaker array and gathering data with adults (Lewald & Ehrenstein, 1998). Because HRTFs were not those of individual participants, we might find auditory localisation to have larger biases than with a speaker array or individually-tailored HRTFs. However, because the experiment included feedback (see below), participants also had an opportunity to recalibrate for such biases. In our analysis, we checked for biases as well as precision.

#### 2.1.4. Procedure

Children were told that they would play a virtual game of hide-and-seek. The headset and headphones were adjusted to be comfortable and ensure clear perception. Participants were introduced to Piccolina, an animated character who would later hide from view. The researchers checked participants could see the entire virtual wall, which Piccolina would hide behind. At this point, the procedure went in three phases: Demonstration, Warmup, and Data Collection (Figure 2).

**Figure 2.**
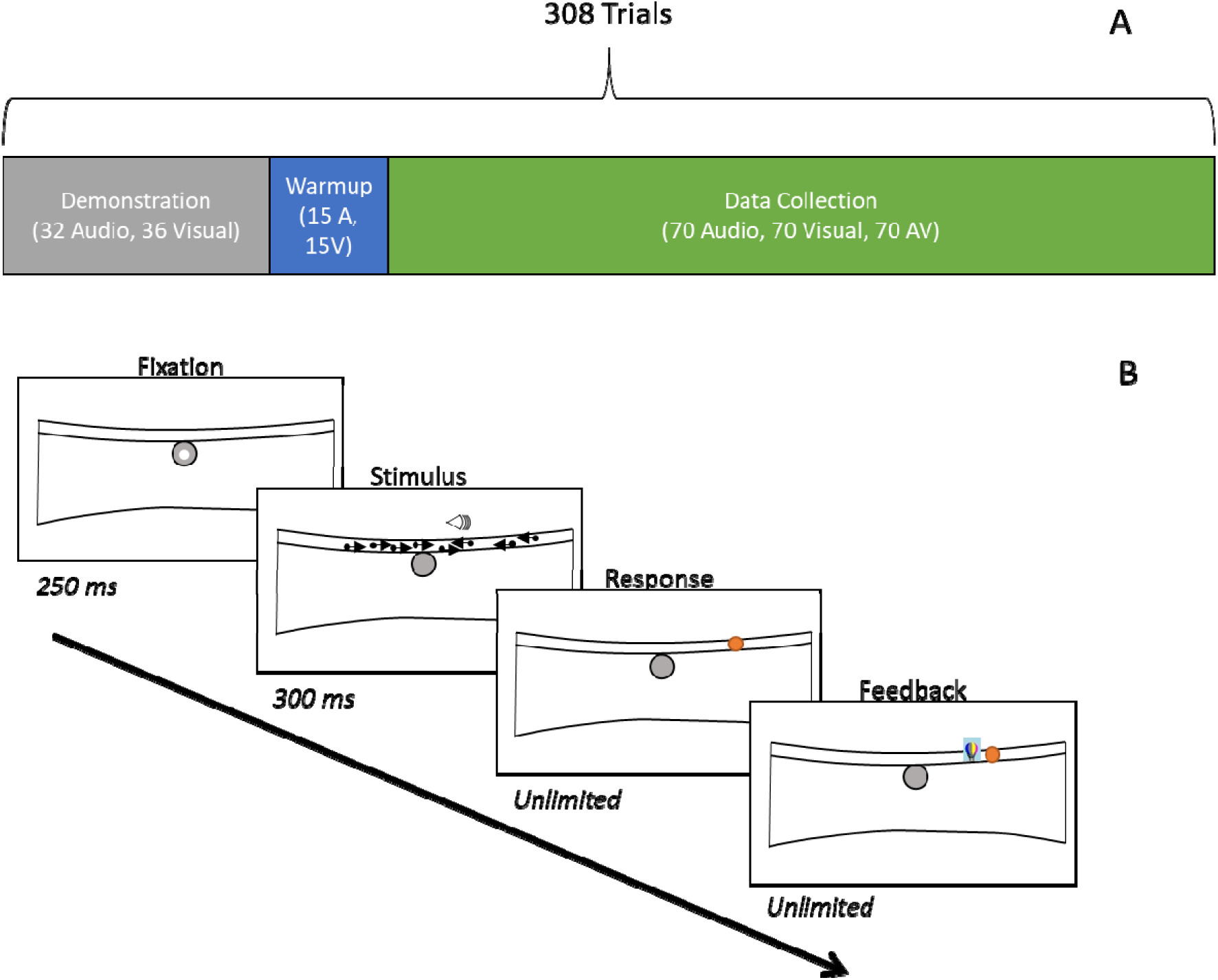
Order (A) and timing (B) of the trials. For the data collection trials, after fixating for 250ms, participants would receive a stimulus that contained audio spatial information, visual spatial information (64 dots moving briefly to a common location), or both. They would then make a response. If there was only one cue presented (audio or visual), they would then receive feedback, which remained visible until they pressed a button to begin the next trial.

##### 2.1.4.1. Demonstration

A gray circle was present in the center of the screen, and a small white dot indicated the central visual point of the headset, demonstrating participants’ fixation (see Figure 1B). Participants were told they could control the white dot by moving their head, and instructed to keep it within the gray circle. If participants moved their heads during the experiment, the task would pause until the white dot was back in the gray circle. This ensured that the visual field remained the same between participants and trials.

Volume was tested. Then children were told they would see dots moving towards Piccolina, as if she held a large magnet. This was demonstrated on-screen, with visual stimuli moving from both directions towards Piccolina, centred on the white wall. The first demonstration of the visual cue was extended, allowing participants to clearly see the dots move all the way to Piccolina. Participants were then told that this was ‘a little bit easy’, and that they would see something a bit shorter. Children were then exposed to a second visual demonstration, showing the dots as they would appear during the task. Children were shown both demonstrations again to consolidate how the cue could be used. Piccolina then moved around the wall, moving in steps towards the far right, then far left, and back to the center-point (32 exposures). At each step, the visual stimuli were demonstrated moving towards her location. Children were instructed that this was how the dots would appear during the task, and could be used as a clue to identify her location. Piccolina then moved around the wall again, and this time children were exposed to the audio stimuli at each of the same 32 positions in the same order. Children were instructed that the sound ‘following Piccolina’ could also be used as a clue to identify her location.

##### 2.1.4.2. Warmup

Piccolina then hid from view behind the top of the wall, and a single-cue trial was given (see Figure 2-B). Children controlled a red dot using a computer mouse to report their location estimate, and a mouse-click to make their judgement. This continued for 30 trials trials, 15 visual-only and 15 audio-only, evenly spaced across the response area. Feedback was given on all of these trials. These data are not used in the analysis.

##### 2.1.4.3. Data Collection

There were 70 evenly-spaced targets for this phase. Each target location was tested once with just an audio stimulus, once with just a visual stimulus, and once with both. Figure 2-B gives the timing of each trial. The order of targets was randomized but the trial type always went audio-only, visual-only, and then both, repeating. Feedback was only given on trials where one cue or the other was presented (not both).

### 2.2. Results and Discussion

Results consistently pointed towards a bimodal precision advantage and thus to cue combination. Participants were excluded if they did not show a significant correlation between target and response for the audio-only trials, the visual-only trials, and the audio-visual trials (N=4 for Experiment 1a). We also focused first on the children under 10 years old. The main outcome measure was the variable error (VE). This was calculated within each participant and trial type (audio, visual, both) as the variance of the signed error, yielding three numbers for each participant. This measure represents the remaining noise after parsing away the mean signed error (bias), with a lower score corresponding to higher precision (better). These VE measures were heavily skewed in many cases, such as a skewness of 2.21 for the audio-visual VE measures, so they were examined with a sign-rank analysis. (This also allows us not to be concerned with potential differences when using standard deviation, variance, or precision; they all give the same rank differences.)

The outcomes of these analyses are reported in Table 1, while the data are shown in Figure 3. The analyses found significant within-subjects differences between VE with bimodal audio-visual cues and each participant’s best single cue in Experiment 1a (Table 1). The bimodal trials showed lower VE (average 78% of best single cue), pointing towards the presence of a bimodal precision advantage. This finding is generally considered to be the most crucial and strongest evidence of cue combination (Ernst et al., 2016) as it demonstrates that the additional (less precise) cue is being used in some way to improve the precision of estimates. Further, we did not find a difference between the audio-visual performance and the prediction of optimal performance (Table 1). The optimal prediction is found by adding the precision from the two cues together (Ernst & Banks, 2002). This means that participants not only showed a bimodal precision effect, but that they further showed an effect that we cannot differentiate from the optimal use of the two cues together.

**Table 1.** Outcome measures for Experiments 1a and 1b (sign rank tests). BS = Best Single; AV = Audio-Visual; O = Optimal.

| <u>Group</u> | <u>Measure</u> | <u>Comparison</u> | <u>Direction</u> | <u>p</u> | <u>N</u> |
| --- | --- | --- | --- | --- | --- |
| 1a: <10 YO | VE | Best Single | BS>AV | .005 | 25 |
| 1a: <10 YO | VE | Optimal | O=AV | .979 | 25 |
| 1b: <10 YO | VE | Best Single | BS>AV | <.001 | 30 |
| 1b: <10 YO | VE | Optimal | O=AV | .909 | 30 |
| 7 < Age < 8 | VE | Best Single | BS>AV | .027 | 9 |
| 8 < Age < 9 | VE | Best Single | BS>AV | .006 | 20 |
| 9 < Age < 10 | VE | Best Single | BS>AV | .003 | 26 |
| 10 < Age | VE | Best Single | BS>AV | .017 | 13 |

Experiment 1b was performed to see if these findings would replicate. They did (Table 1). Four children were also excluded on the same criteria (remaining N=30). There was a significantly lower VE for the bimodal trials compared to the same participant’s best single cue (on average, 73%). There was no difference between the audio-visual performance and the predicted optimal performance. This suggests that these findings are reliable and replicable.

For further analysis by age group, the data from the two experiments were pooled together for a larger sample. Each year-long age group showed a significant bimodal precision advantage in terms of VE (Table 1, Figure 3). On average, the bimodal VE was 82%, 74%, 73% and 76% of the best single cue VE, for each age group in order. This suggests that the VE effect in each experiment is not driven just by the upper part of the age range, but that this task is solvable even at 7 years old.

**Figure 3.**
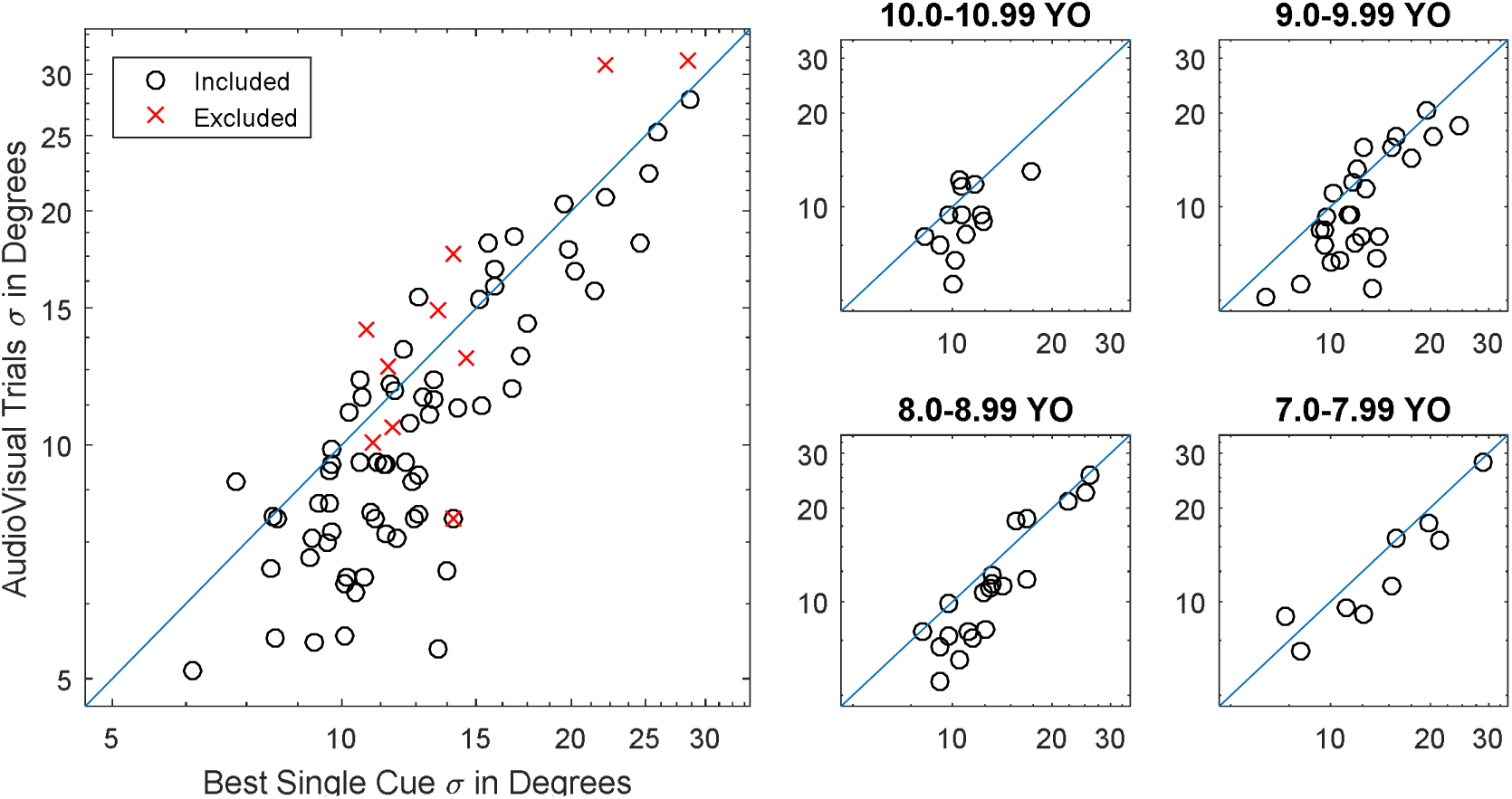
Comparing the standard deviation of responses away from the targets with the best single cue (X axis) versus the two cues together (Y axis). The reference line in blue is where they are equal. Data points represent individual children. Points below the line indicate a bimodal precision advantage. Points nearer the left represent a better use of at least one cue. The left panel shows all children tested, with the right panels breaking this apart by age.

The bimodal precision advantage was also present and significant when looking at all data collected with children under 10 years old (see Figure 3), including the 8 children that were excluded, N = 63, p < .001, showing that this effect does not depend on our exclusion criteria.

Biases were centered near zero (see Figure 4) and audio-only biases were comparable in magnitude with previous reports of typical adult biases for audio-spatial stimuli (Lewald & Ehrenstein, 1998). This suggests that our use of HRTFs through headphones was not a major issue. Further, using the VEs, we can estimate the weight that would be optimal to give to each cue during audio-visual trials (weighted by precision, which is 1/VE, as measured in single-cue trials). We can then predict, for each participant, the bias in the audio-visual trials if the two cues are being averaged with the optimal weights. The predicted and observed biases in the audio-visual trials were significantly correlated (Figure 4, right), *r*(108) = .56, *p* < .001. This is also consistent with cue combination.

**Figure 4.**
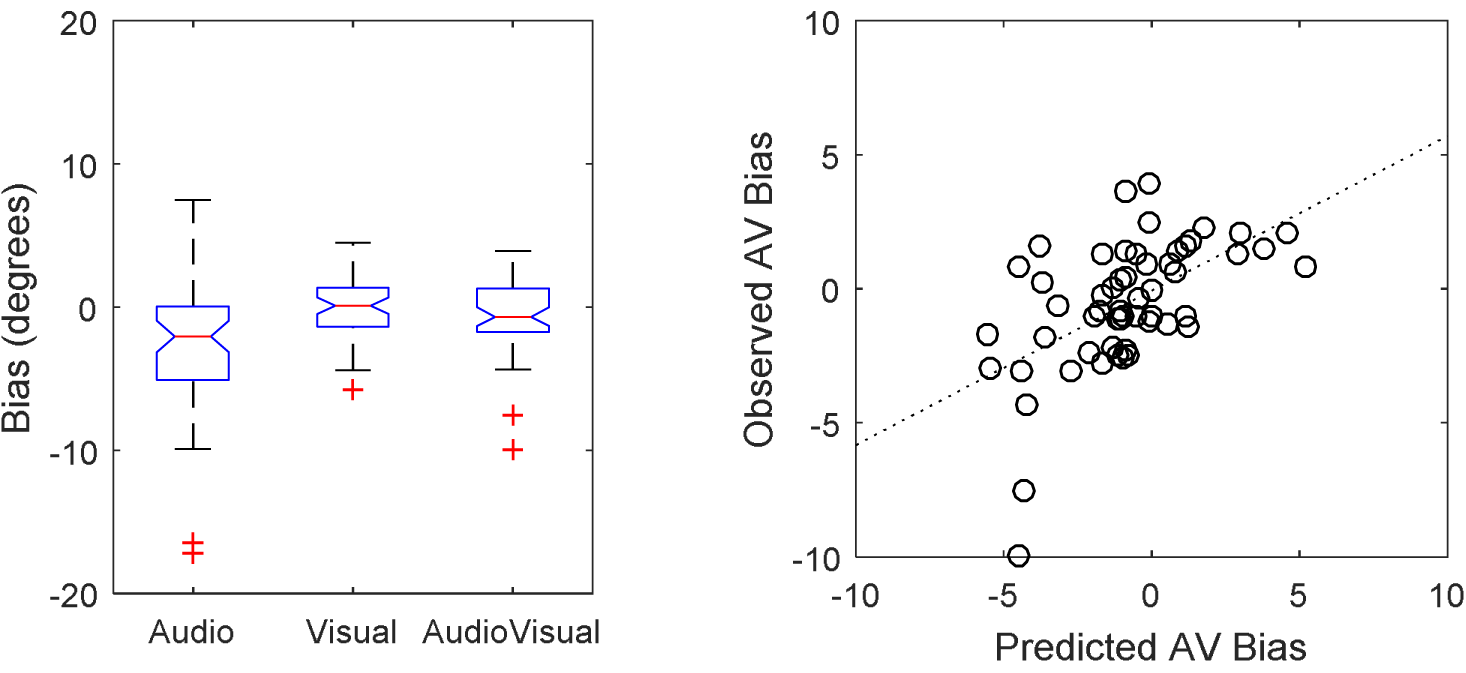
Biases in the task. On the left, a box and whisker plot of observed biases in the included children under ten years of age. Notches represent a 95% confidence interval on the median. The red line is the median. The box spans from the 25^th^ to 75^th^ percentile. The whiskers extend two 1.5x the interquartile range or the furthest data point there. The red crosses are further data outside the whisker range. On the right, the bias in the audio-visual trials was significantly predicted by the optimal combination of the two cues.

To further deal with biases, we also looked at the mean squared error (MSE). This measure includes the bias, creating an overall measure of accuracy, which may be of more practical interest. It was calculated by squaring and then averaging the signed errors. (In contrast, the VE subtracts out the mean signed error.) This bimodal MSE was, on average, 90% and 88% of the best single cue MSE for each respective experiment. This also reached significance in Experiment 1a, p = .019, and Experiment 1b, p = .006, in a sign-rank test between the best single cue trials and the audio-visual trials. This suggests that participants not only increased precision, but also accuracy, with both cues available.

To see how this pattern of performance was reflected in the raw data, we show the full target and response distribution of a selected sample participant with lower audio-visual VE than either single cue VE in Figure 5. Full data are also attached in Supplemental Materials for any further analysis desired by future researchers. Comparison with the identity line shows that bias is relatively minor (especially in the center of the range). The lower variability along the y axis in Audio-Visual than either single cue condition marks the bimodal precision advantage.

**Figure 5.**
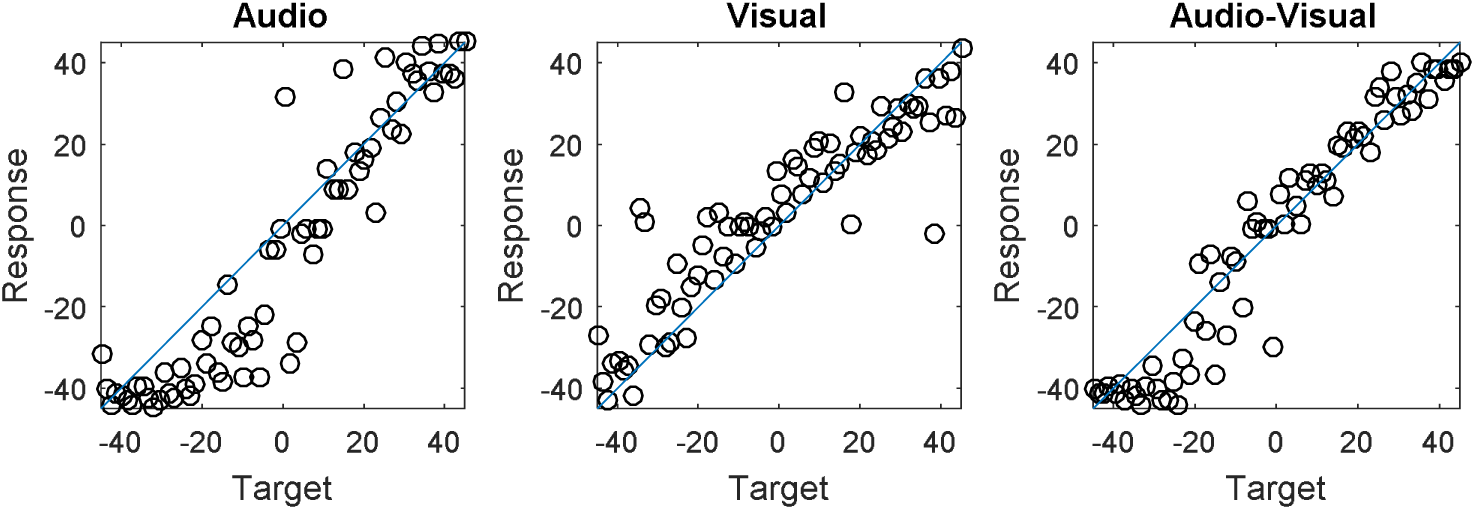
Full data from a selected representative participant. The AV trials tended to be closer to the target (i.e. fall closer to a line from the bottom left to top right), whereas the response to each single cue were less precise. Points represent individual trials.

In summary, the results of both Experiment 1a and 1b suggests that children under 10 years old can combine cues in this task, perhaps even optimally. This is robust to a wide variety of ways of looking at the data as well – various outcome measures, experiments, and subgroups of the age range. This violates the predictions of the Hard Limit hypothesis and instead matches the predictions of the Undiscovered Task hypothesis.

While results from Experiments 1a and 1b already point towards favouring one of the two key hypotheses, matching the standard of evidence set by a previous project (Dekker et al., 2015) requires the effect to be shown independently a third time in a different kind of measure. We chose a model comparison measure as the strongest way of verifying this. If children under 10 years are combining cues, they should not only show a bimodal precision advantage, but a combination model of their data should be preferred over a model that posits the use of a single cue on each trial. Experiment 2 fills this role.

## 3. Experiment 2

Here, we prioritized the ability to carry out model comparison with each individual child’s dataset separately. In this case, we are not primarily testing for a bimodal precision advantage with a null hypothesis of no difference and a vague alternative of some difference (though this effect was also observed). Instead, we created several generative models of the algorithm used to deal with the two sources of information, derived predictions of the bimodal trials from performance on the single-cue trials for each child and each model, and compared how well these predict the actual bimodal data for each child.

This means that we are looking at the best model for each child, not the entire group. This is a challenge with children at this age because they are unlikely to be willing and able to focus on the task for long hours or thousands of trials. Instead of raw trial numbers, we used a visual stimulus with external noise (under the control of the experimenter) rather than internal noise (a function of the participant’s noisy perception). This gains us a significant amount of statistical power for certain comparisons. For example, suppose an audio cue is directly on the target but a visual cue is placed slightly to the right of it. If the visual cue is being used in isolation, we do not just expect noise to be larger than it would be under optimal combination; we expect error to be in a specific direction (rightwards) and a specific magnitude range (the displacement amount, plus or minus an allowance for motor noise). We have a much narrower prediction available than if we just presented two cues at the same location with internal noise, allowing much more power in the model comparison. (See the prior predictive distributions in Figure 7 below for a concrete example.)

We also wanted to look more at the role played by feedback. We did not want to have a condition without feedback as this may simply not be as motivating. However, we could instead manipulate if they received feedback on single-cue trials (like Experiments 1a-b), the audio-visual trials, or both. Feedback on the audio-visual trials might be more helpful than single-cue trials because it makes it the most obvious when cue combination would have led to a more accurate response, and could potentially train a cue combination strategy. On the other hand, feedback on single-cue trials was already very effective in Experiments 1a and 1b. It is also possible that learning relative to both cues at the same time is difficult for young participants. In any case, an effect where differences in feedback lead to different rates of cue combination behaviour would support the general idea that feedback is an important variable for predicting the combination of cues.

Further, we wanted to see if participants could stretch beyond the predictions shown by the experiment above. The full remit of Bayesian Decision Theory predicts not just a precision advantage for combination of cues, but also that this should persist when the relative reliability of the two cues changes – even trial to trial. To test this, we allowed the precision of the visual cue to vary randomly within a range around the running estimate of the audio cue’s reliability for each participant. This also helps ensure that the reliabilities will tend to be reasonably matched for most trials.

### 3.1. Method

#### 3.1.1. Participants

Eighty-three participants were tested, some in the lab (N=22) and the rest in a local school. Of these, 16 were excluded for failing to show a significant correlation between targets and responses in the audio-only, visual-only, or audio-visual trials, leaving 67 participants. Ages ranged from 7 years and 0 months to 9 years and 11 months (17, 27, and 23 respectively in each year range). The ages were distributed evenly among the three conditions, with each having an average participant age of 8 years and 4 months. This study was also approved by the Psychology Department Ethics Sub-Committee at Durham University (reference: 16–07 Multisensory Learning). Parents gave written informed consent.

#### 3.1.2. Apparatus and Stimuli

The apparatus was the same for participants tested in the lab and in schools in terms of display equipment, headphones, and environment, but in the lab setting we gave them a virtual laser pointer as a response device instead of a mouse. In both cases, the audio stimulus was the same as Experiments 1a and 1b. The virtual environment was also the same as the previous experiments, except for the visual stimulus.

The visual stimulus was a display of a normal distribution placed over the wall (see Figure 6). The targets were generated by actually drawing from this distribution and re-drawing if this fell off the ends of the wall, effectively making the visual display a depiction of a likelihood function. This means that the visual cue went from having internal noise to external noise for this experiment. Further, crucially, the variance of the visual cue varied trial-to-trial from +25% to −25% of the running estimate of their audio variance. This was signalled by drawing the visual cue wider or narrower on each trial. This cue’s characteristics challenged participants to not only combine the cues, but to do so with varying weights across the experimental session.

**Figure 6.**
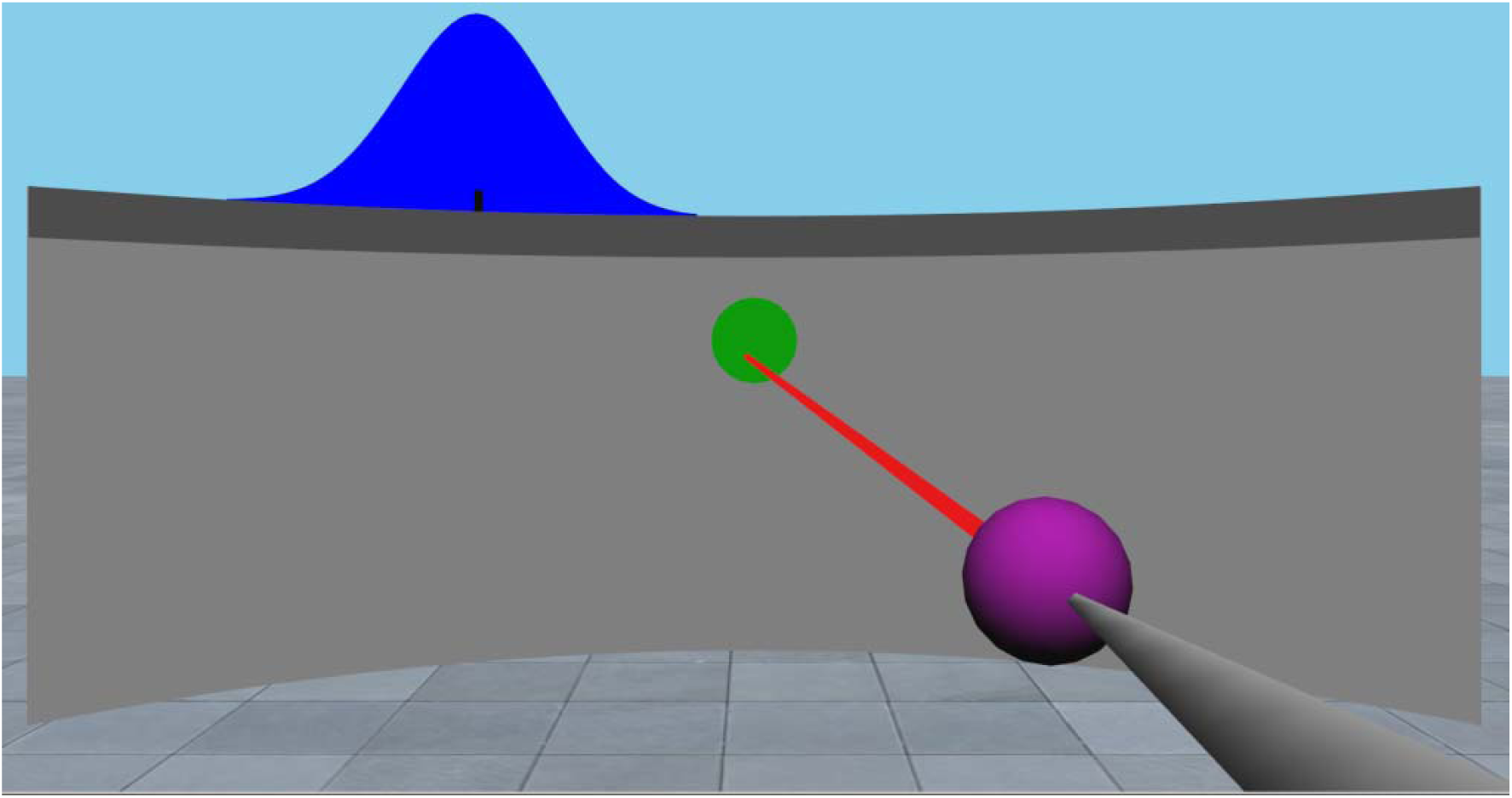
Screenshot of the visual stimulus and the alternative response device used for in-lab testing. The blue curve on top of the wall can be interpreted directly as a likelihood function. The target is most likely to appear directly under its center, becoming less likely to the left and the right, exactly in proportion with its depicted height. Crucially, the width and placement of this cue varied trial to trial.

#### 3.1.3. Procedure

Procedure for this experiment was very similar to Experiment 1a and 1b but had two notable differences. First, the feedback varied by condition between subjects: all trials (feedback on audio, visual, and audio-visual), single-cue only trials (feedback on audio, visual), or audio-visual trials. To make the feedback amount similar in the single-cue feedback condition and the audio-visual feedback condition, the participants in the audio-visual feedback condition experienced twice the number of audio-visual trials (rotating audio, audio-visual, visual, audio-visual).

Second, the demonstrations were altered. The audio demonstration was shortened to 16 trials. The visual demonstration was altered to be three stages of 16 draws each. The first draw was from a central, medium-variance visual cue. The second was from a leftwards large-variance visual cue. The third was from a rightwards low-variance visual cue. This was intended to introduce the idea that the reliability of the visual cue could change and that the highest point on the visual cue indicated the most likely target.

#### 3.1.4. Models

Here we give a brief explanation of each model. Appendix A specifies the details of the model fitting and the priors given on each model. Figure 7 shows the prior predictive (the probability distribution of *a priori* predicted responses) for each of the six models given some example trial parameters and single-cue trial performance parameters.

**Figure 7.**
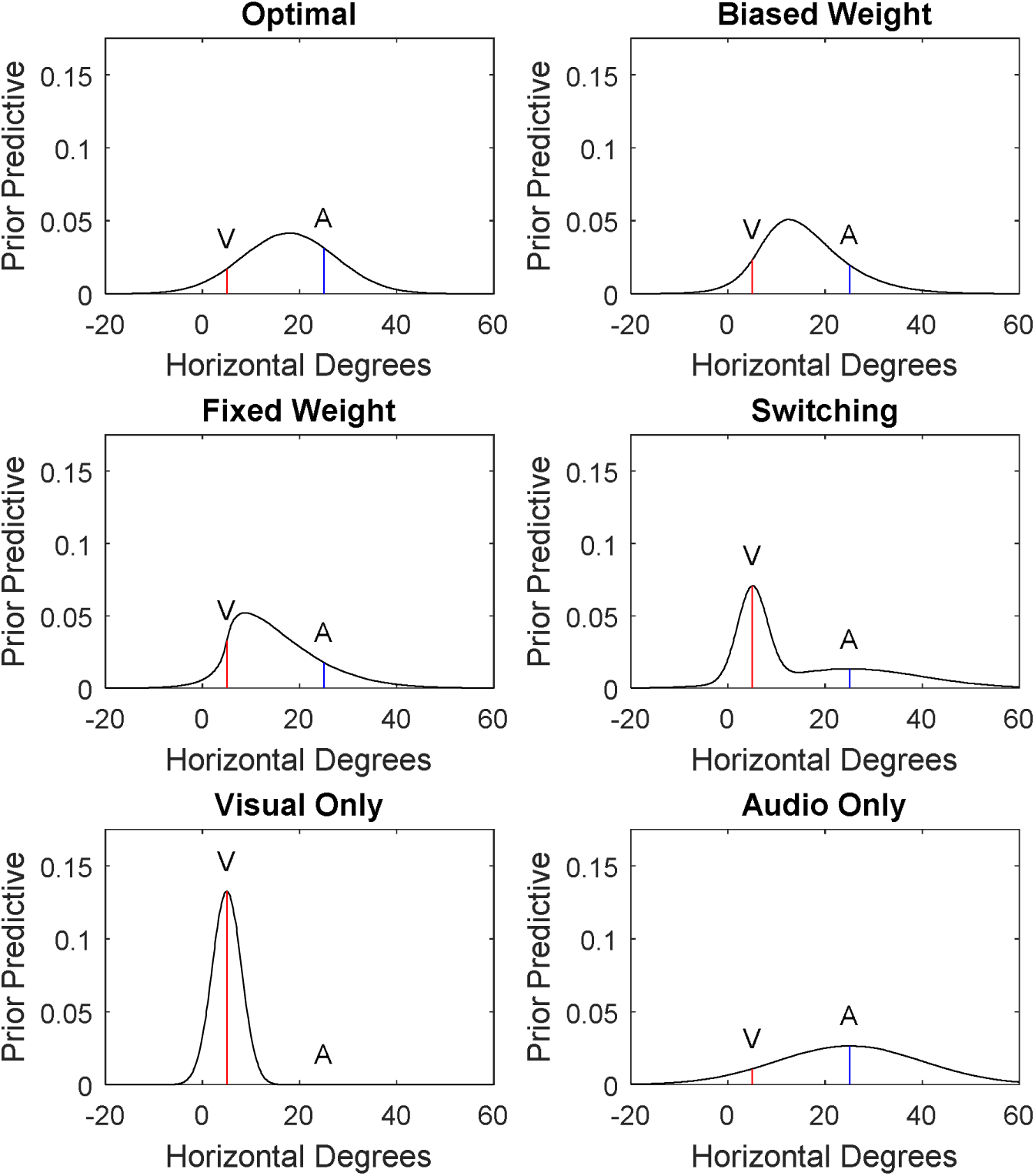
Prior predictive probability distribution for each model given an audio cue at +25 degrees (marked A with a blue line), a visual cue centered at +5 degrees (marked V with a red line), a 3 degree standard deviation in their responses when pointing to the center of the visual cue, a 15 degree standard deviation in their audio perception, and the visual cue subject to an additional 20 degrees of external noise (induced by the experimenter by changing the spread of the visual cue; see methods). The models each took in fits from the audio-only and visual-only data and then had to predict how the audio-visual data would be distributed. Models were assessed on the basis of how well these predictions matched the actual audio-visual data. For the visual-only, we can make very sharp predictions because the placement of the visual cue was under experimenter control (external noise); they are expected to just point near its center. In contrast, for the audio-only, predictions are wider since they include the noise in audio perception (internal noise). The rest are various ways of mixing the two, either switching between them from trial to trial or averaging (combining) them with different weights.

##### 3.1.4.1. Optimal combination

In this model, the two estimates are weighted by their relative precisions and averaged. This represents the best possible use of the two cues for maximising precision around the correct target. Note that under this model, the weights change trial-to-trial based on the varying reliability of the visual cue (c.f. single weighting below). In the example (Figure 7), this leads to a distribution peaked between A and V. It has lower variance that the audio-only model and also lower variance than the visual cue’s true external noise.

##### 3.1.4.2. Biased weighting

This is much like optimal combination, except the weights are corrupted by multiplying the visual cue’s precision by a fixed factor. If this factor is above 1, the cues are combined but the visual cue is given undue weight. This factor was given a relatively narrow prior around 1 since a very strong over-weighting of the visual cue is very similar to just using the visual cue in isolation (and the same for under-weighting and audio). In the example, the distribution is much like the Optimal but shifted more towards the visual cue’s center.

##### 3.1.4.3. Fixed weighting

This is also like the optimal model, but it is unresponsive to the relative precision of the two cues (both across subjects and across trials). A single fixed weight for the visual cue is set between 0 and 1 and applied for all trials. In the example, this leads to a skew towards the visual cue’s center as it frequently receives undue weight.

##### 3.1.4.4. Switching

The participant has a probability of choosing either single cue and just using it alone. They never use both cues on any single trial, but they do use both of the cues on separate trials over the course of the experiment. In the example, this leads to a bimodal distribution.

##### 3.1.4.5. Visual only

The participant just uses the visual cue. Their responses should be similar in distribution as when they only have the visual cue available: centered on the visual cue’s center and unresponsive to variation in the placement of the audio cue. This is actually the prior predictive with the least variance in the example, since the visual cue had external noise; this allows us to make a very narrow prediction about responses.

##### 3.1.4.6. Audio only

This is exactly like the visual-only model but instead just relies on the audio cue. This leads to a distribution centered on the audio cue, with a wider variance than the optimal model.

### 3.2. Results and Discussion

Each participant’s data was analysed separately through a Markov Chain Monte Carlo procedure (4 chains, 25,000 samples, 5,000 used as burnin). The main outcome is which model fits each participant best in a Bayes factor sense (i.e. probability of the model given the data). Since these models all approach the audio-only and visual-only data the same way (a linear regression to the targets), this reduces to asking which can predict each participant’s audio-visual data the best from their audio-only and visual-only data. For example, under the parameters used to generate Figure 7, a response at +5 degrees would be best predicted by the visual-only model, but a further response at +25 degrees would be poorly predicted by the visual-only model and would start to favour the switching model.

Most participants (34) had a decisive fit, with the best fitting model having a Bayes factor of at least 10 times above the second-best. Figure 8 gives the counts of the best fitting models by feedback condition. Overall, results suggest that the majority of participants were combining cues (in total 65% of children combined) either optimally (33%) or with biased weights (30%), and with one child best fitted by the fixed weight model. Strategies relying on one cue on each trial (i.e. no combination) were rarer, with 19% switching between trials and 16% just using the visual cue. While this highlights heterogeneity in this age range, it still speaks strongly to the idea that cue combination in this task can be done by children under 10 years.

**Figure 8.**
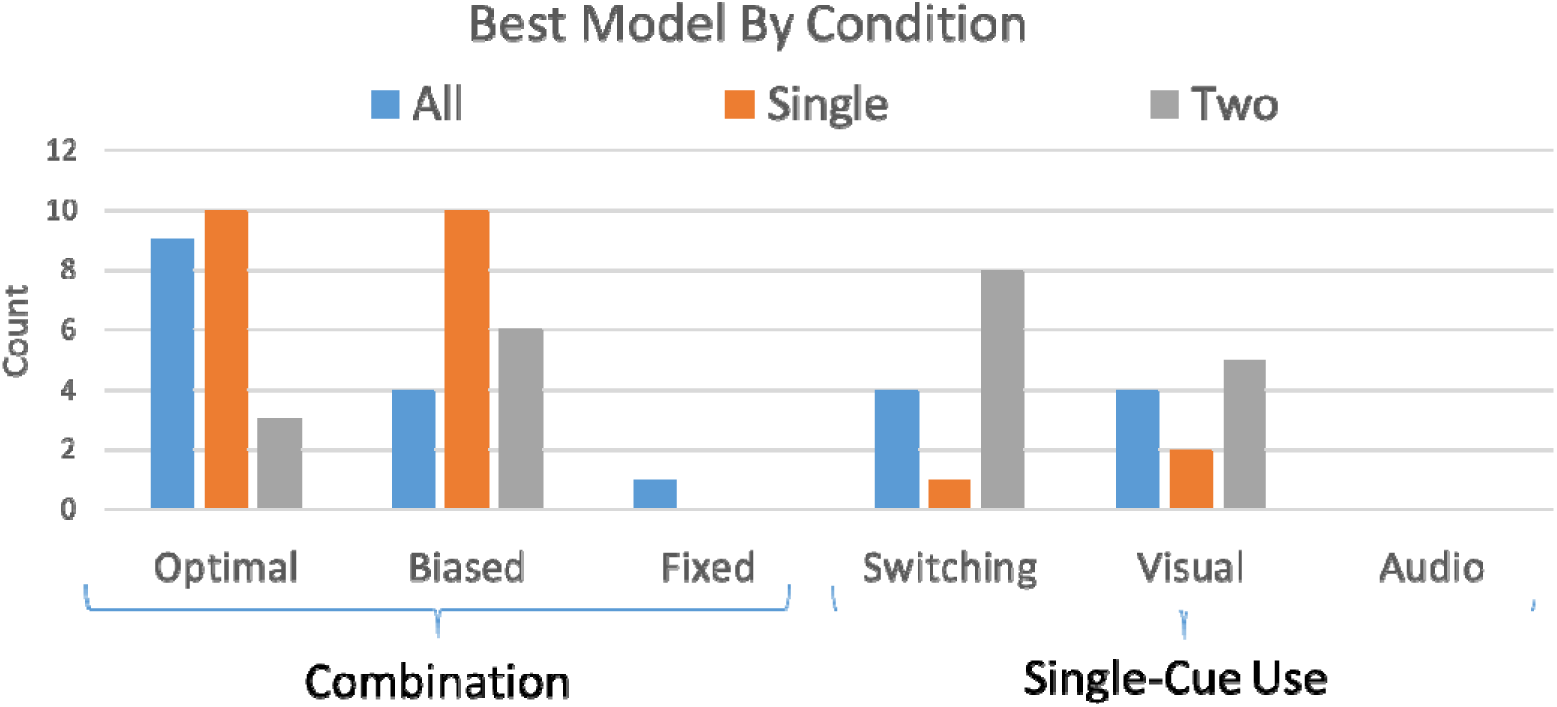
The count of participants best fit by each participant in each condition. The majority of children were best fit by a combination model (65%), especially in the single-cue feedback condition (88%). Biased weighting of the two cues was a very common non-optimal strategy, as were switching and relying solely on the visual cue. However, few children failed to respond trial-to-trial to the varying cue weights (fixed weight) or used the audio cue exclusively.

To test the effect of conditions with a chi-square, the counts of the three combination models (optimal, biased, fixed) were collapsed, as were the counts of the single-cue use models (switching, visual-only, audio-only), because many individual cells had zero data points within them. A significant association between feedback condition and model category was found, N = 67, *X*^2^(2) = 10.38, p = 0.006, suggesting that the different levels of feedback led to different rates of cue combination. The condition with the most models in the combination category was the single-cue feedback condition (88% combining optimally or sub-optimally). This was lower in the conditions with feedback on all trials (64%), followed by feedback only on audio-visual trials (41%). This is still consistent with the hypothesis that this age range is capable of combining cues in this variation of the task. However, it does suggest that feedback on single-cue trials is generally more helpful in aiding cue combination than feedback on dual-cue trials. This difference emphasizes a role for the feedback in predicting cue combination.

Another notable finding is that we found only one child best fit by the fixed weight model and none best fit by the audio-only model. This might indicate that for children at this age, the major issue is not (a) being incapable of having their responses change based on changing relative reliability or (b) allowing audio capture for a spatial judgement. Instead, the major issues seem to be (1) learning the correct weights for the two cues, (2) deciding to integrate them both rather than switch back and forth between, and (3) using an audio cue at all for a spatial judgement – in a sense, the most extreme example of incorrect weights. In addition, the preference for the visual-only model over the audio-only model converges with a previous project (Gori et al., 2012), so we now have consistent evidence that visual capture is a major issue for spatial judgements with audio-visual cues in development.

We also did two things to make sure that the models were working as intended. First, we generated 100 datasets of 210 trials from each model and ran them through the comparison. These simulations used a gamma distribution of noise levels in each cue that was similar to the actual child data. We found that the method recovered the generating model on 96.8% of datasets. This confirms that the models are differentiable and that the analysis method is a reliable way to identify the correct model among these. Second, we reasoned that there should be a bimodal precision advantage as seen in the previous experiments 1a and 1b if the model fits are correct. We found indeed that there was, with the bimodal trials showing lower variable error than the best single cue trials in a sign-rank test, N = 67, p < .001.

## 4. General Discussion

All three experiments provided results converging on cue combination in this task in children over 7 years and under 10 years old. This can be interpreted alongside another report about the combination of motion and disparity cues to depth (Dekker et al., 2015). That study provided three independent analyses that converged on children in this same age range failing to combine cues, but instead doing so over the age of 10.5 years. This further sits within a larger literature where there have been 25 separate failures to find a bimodal precision advantage under the age of 10 years old. Taken together, our current data and previous results provide convergent evidence that cue combination emerges at different time points for different tasks. As a whole, this is compelling evidence against the idea that cue combination emerges in a unified fashion in development, instead strongly suggesting that children under 10 years old can combine cues – they just need the right task.

The model comparison results in Experiment 2 also clarify which of the proposed alternative strategies are most likely to fit performance when it is not best fit by optimal cue combination. The most common alternative is combination with the incorrect weights. We also saw many participants discarding the audio cue entirely, converging with a previous report where children in the same age range showed a similar effect (Gori et al., 2012). This suggests that it remains tempting through middle childhood to over-rely on vision for spatial judgements, sometimes even to an extreme where audition has no effect at all.

We also found further evidence for an effect of feedback on cue combination behaviour in Experiment 2, but not the simple one that might be expected. The simplest story to tell is that children between 7–10 years combine cues with feedback, like our Experiments 1a-b, but do not combine cues without feedback, like a previous audio-visual study in the same age range (Gori et al., 2012) which found strong visual capture. But as our Experiment 2 shows, the full results are more nuanced: some appreciable level of cue combination behaviour was seen in all three feedback conditions, but the condition with single-cue only feedback showed the highest rate (88%) and the condition with dual-cue only feedback showed the lowest (41%). This suggests instead that single-cue feedback effectively promotes cue combination behaviour, changing the rate of cue combination observed, but not acting strictly as a binary switch.

Finally, only one child was best fit by a model with fixed weights in Experiment 2. Under this model, children do not adapt on a trial-by-trial basis to the varying reliability of the visual stimulus. They instead set one weight for the visual information and apply it regardless of the varying reliability. The result that this model was rarely selected indicates that children, once they are no longer selecting cues, quickly come to understand that the weights given to each modality must vary based on the nature of the stimulus presented (Maloney & Mamassian, 2009).

The primary finding is a key guide to the development of future theory. At the moment, papers that model the acquisition of cue combination tend to discuss emergence as a unified whole (Daee et al., 2014; Weisswange et al., 2011). Current models also do not (at least explicitly) differentiate various tasks in ways that predict their different courses of emergence. Future models should take on this challenge and attempt to understand why the same computations might emerge at different developmental points in different tasks. Further understanding of the order of emergence will let us see the crucial contours of what makes an ‘easy’ or ‘hard’ learning problem, thus gaining insight into the learning process.

### 4.1. Favoured Explanations

This finding opens up a new theoretical depth in our understanding. With this result in hand, it is now sensible to ask what exactly makes this task different from the large body of attempts to find a similar bimodal precision advantage in this age range. In this section, we present and critically evaluate the two explanations that seem most likely to us. In the section after, we argue why some competing explanations are less compelling.

#### 4.1.1. Feedback Adjusting Perceived Relative Reliability

Under this hypothesis, participants come into any given cue combination task in the 7–10 year old range equipped with the (optimal) reliability-weighted-averaging algorithm. However, they lack correct estimates of the relative reliability of the two cues. Without this, they default to a switching strategy, default to a cue selection strategy, or combine the cues with incorrect weights. No bimodal precision advantage obtains. In contrast, in our experiments, the feedback allows them to estimate the correct weights with reasonable accuracy and then create a bimodal precision advantage through cue combination.

The question then is why natural experience would not give children under 10 years good estimates of relative reliability. Here we think there is at least one plausible answer. The problem of estimating the reliability of a wide variety of audio and visual stimuli, in terms of their localization, may be too overwhelming for various reasons. It might place memory demands that are too high or require them to develop a fuller internal model of their own perception than they currently have. The present task, in contrast, requires them only to derive an estimate of the reliability of the subset of audio and visual cues that we presented. It seems plausible that this smaller problem is tractable but the larger issue of a good general model of own uncertainty is not.

It is also worth noting that a previous study which directly assessed weights in an audio-visual task found that they are frequently far from the optimal weights in this age range (Gori et al., 2012), lending plausibility to the explanation above. Given the results by condition in Experiment 2, we also expect that feedback must play some sort of role in the correct explanation.

#### 4.1.2. Feedback Reducing Bias

Under this hypothesis, biases in perception prevent cue combination and the feedback acts against that effect. Suppose we present two cues that a target is at some specific point. The participant has a particular set of relative biases. When both Cue A and Cue B signal the same point, the participant tends to perceive Cue B as signalling something +10 higher than Cue A. If the standard deviation of perceptions is well less than 10 for both cues, the participant will typically perceive the two cues to be signalling irreconcilable states of the world. Under these conditions, an optimal computation rejects the idea that the two cues have the same cause and simply uses the more reliable of the two cues in isolation (Knill, 2007; Shams & Beierholm, 2010). This kind of filtering makes perception more robust to unexpected issues, but, in this hypothetical, does so at the cost of a bimodal precision advantage.

The feedback would most likely help correct such an issue. It would provide an objective way to see that perceptions from one cue or the other (or both) are not just noisy, but on average biased in one direction or the other and thus should be corrected. This also fits well with theory regarding cross-calibration (Burr & Gori, 2012), as the single-cue feedback can serve for this calibration. This also generally fits with the results of Experiment 2, which suggests some role for feedback in the current findings. However, we are not aware of any documentation that children in this age range actually have such severe biases.

As above, it seems plausible that natural experience would not be sufficient to help children find and correct relative biases for a wide range of possible stimuli in the audio and visual domain, but that the focused task of correcting biases on these specific stimuli might be within reach.

Of course, this explanation is not exclusive with the one above – children at 7–10 may both have biases and incorrect weights that need to be adjusted for a bimodal precision advantage. The feedback here could fill both roles.

If we accept these explanations, it shifts some of the texture of what cue combination learning models need to explain. At the moment, models take on the general idea of audio-visual cue combination as essentially one task where an audio stimulus in horizontal location has the same reliability as all further possible audio stimuli in the same place (Daee et al., 2014; Weisswange et al., 2011). The estimated reliability parameter for each modality can adapt over long times (thousands of trials), but there are not, for example, separate reliability estimates for a dark room versus a brightly lit room. The way that we are discussing it here, learning the relative reliability of these specific audio-visual stimuli is not the same as developing an overall model of how reliable one’s own auditory and visual system are under different circumstances. Current models do not make it obvious how that kind of segmentation would occur in cognition.

### 4.2. Unlikely Explanations

In this section, we argue why several additional tempting options for explaining the current results are unlikely.

#### 4.2.1. Exposure Frequency

At the moment, current models do have one implicit aspect that could differentiate some pairs of cues (Daee et al., 2014; Weisswange et al., 2011). Specifically, they assume that learning is mainly a function of the number of joint exposures to the cues, plus some noise. Therefore, one tempting possibility is that different cue pairs occur with different frequencies in naturalistic settings and that children learn to combine the most common ones first. In a broad sense, we do know from deprived rearing tasks that this is a relevant variable – for example, cats raised in deprivation of co-varying spatial sounds and lights will fail to show audio-visual localization effects that are consistent with cue combination (Xu, Yu, Rowland, Stanford, & Stein, 2014).

However, this explanation faces some difficulties. First, it is not clear that audio-visual localization is something that children can practice more frequently than visual depth (Dekker et al., 2015), and it fails to explain why the other audio-visual study (Gori et al., 2012) did not find a bimodal precision advantage. Second, modelling the acquisition of cue combination as purely a function of cue exposure runs aground of a recent finding in adults (Negen, Wen, Thaler, & Nardini, 2017). In a 5-hour task, adults were able to combine a new audio cue to depth with a familiar but noisy visual cue to depth. Therefore, something besides experience is also a strong predictor of learning to combine cues; otherwise, children would combine things like visual depth cues well before 10 years of age. Perhaps then, it is best to think of a certain level of exposure frequency as being necessary but not sufficient to induce cue combination in a specific task.

#### 4.2.2. Better task engagement in our task than previous tasks

Experiment 2’s results suggest that this is, at least, not the entire explanation for results here. Participants still varied in how likely they were to combine cues across conditions that all had feedback, all had a social entity, and all took place in virtual reality. Large differences in the level of engagement seem unlikely. It also seems unlikely that all of the previous projects failed to reach a high level of task engagement. Like exposure frequency, we interpret this as being necessary but not sufficient.

#### 4.2.3. Feedback Providing Participants with a Look-Up Table

This possibility has been discussed by previous theory at length (Maloney & Mamassian, 2009). Under this hypothesis, training allows participants to associate each specific pairing of single-cue perceptions with a specific response. In theory, the observer can then make accurate responses by recalling the best response to each stimulus pairing from past experience (i.e. using a “look-up table”), and could even show a bimodal precision advantage with this algorithm. However, this is not a possibility for the specific design here: in Experiments 1a, 1b, and the “Single” condition of Experiment 2, we gave no feedback on trials where they had both cues available, yet participants still showed evidence of cue combination. Even beyond that, participants who were only given feedback on trials with both cues actually became less likely to combine them.

#### 4.2.4. Feedback Teaching Participants the Correct Algorithm

Along similar lines, a model learner could keep track of the outcomes of their perceptual decisions and see which of several algorithms leads to the best performance. In other words, if you look over the models we compared in Experiment 2, one might imagine a child doing the same kind of thing in their own mind – but instead of an external experimenter trying to decipher their preferred strategy, the participant is trying to decipher the strategy that will lead to the best performance. This is again a poor fit for the current results for the same reason as above, the lack of feedback with both cues available. Most participants would not have anything to fit this against, and the ones who did were less likely to combine cues.

### 4.3. Related Issues

Given recent findings in adults, we should also point out a related quandary about development that is also very open. While this study is the first to show cue combination in children under 10 years, it is perhaps not actually unique in showing that the combination of some cues follows a radically different developmental path from visual depth cues. Specifically, even adults don’t combine certain cues to motion (e.g., Soyka, de Winkel, Barnett-Cowan, Groen, & Bülthoff, 2011), or at least vary in whether or not they combine them (e.g., de Winkel et al., 2013). This is somewhat difficult to explain under current models of multisensory learning (Daee et al., 2014; Weisswange et al., 2011), which should always converge on cue combination for any two common cues to a common judgement. Thus, a truly accurate model of how cue combination is acquired must explain why this task is learned earlier, unreinforced visual depth is learned later, and things like angular self-motion might not necessarily be learned at all.

Another important discussion point is how the studies of the full realm of Bayesian Decision Theory should proceed for development. This theory, which suggests that perception is fundamentally well-represented by a model of internal Bayesian statistics, creates a wide range of predictions (though c.f. Rahnev & Denison, 2017). We have taken a step in this direction by not only showing a bimodal precision advantage, but showing that a model where children re-weight cues trial-to-trial fits better more frequently than one with a single fixed weight. This shows some of the re-weighting predictions to be sensible in this age range, at least for the right task. Some additional points to consider are the question of when and how children learn to learn prior distributions, learn to maximize gain functions rationally, learn to calibrate confidence, and acquire a causal inference filter. Compared to cue combination, relatively little is known about each of these. Understanding that full breadth is important for a complete theory of multisensory perceptual development.

### 4.4. Towards Applications in Autism, Robotics, and Sensory Augmentation

This task and finding could be used in the future as a comparison point to work towards a deeper understanding of sensory atypicalities such as those associated with autism or schizophrenia. Various theories for various conditions have posited specific differences involving the integration of information from different modalities (e.g. Baum, Stevenson, & Wallace, 2015), while others have focused on alternatives like the learning of prior information (Lawson, Mathys, & Rees, 2017). The present study has been able to take a closer look at typical development and discover a multisensory integration capacity that was not known before. It could be informative to explore if the bimodal precision advantage measured here is associated with neurodiversity in any systematic way. For example, the feedback might be used in different ways by different populations.

In terms of robotics, our results point towards an interesting suggestion for self-organization. We posit that the children in our task were not generally capable of solving any arbitrary audio-visual cue combination problem (Gori et al., 2012), but that the feedback given here allowed them to create a temporary working solution for a narrow range of audio-visual stimuli (specifically those used in the task). This might be part of the path forward for a robot attempting to adapt to different environments as well (Lake et al., 2017). It might be most effective, or at least most human-like, if a robot were to develop smaller solutions to more restricted and common cue combination problems and then to generalize much later from that set of smaller solutions.

In terms of sensory augmentation, two studies have been done that stand in stark contrast. One found that adults can integrate augmented sensory skills with their typical repertoire (Negen et al., 2017), but the other found that they did not (Goeke et al., 2016). The study with feedback showed integration, but not the study without feedback. The present study shows that in young participants, feedback can be an important variable. This suggests an overall pattern that remains important into adulthood, with a strong emphasis on frequent feedback in order to help participants learn how to integrate a sensory skill. It may be that single-cue feedback is particularly effective for adults learning to integrate a new sensory skill as well.

### 4.5. Conclusion

Across three independent experiments, we found evidence that children between 7 and 10 years old can combine audio-visual cues to location in a way that is more precise than either single cue alone. This establishes that cue combination is possible in some tasks under 10 years old. Comparing with previous research, this further establishes that cue combination emerges at different points in development for different tasks. We propose that this is most likely to be due to the feedback given during the task, rather than the other details of the task chosen. This interpretation suggests specifically that children between 7–10 years are able to combine cues with some intensive experience to calibrate the two cues (correcting biases and/or assessing relative reliability), but will not do the same if forced to rely on their natural experience with them. Current models of the developmental process do not account for this and should be extended to create a better theory of how cue combination develops.

## 5. Acknowledgements

This work was supported by Economic and Social Research Council of the United Kingdom ES/N01846X/1.

## Appendix A

In this appendix, we give the full parameters, priors, and other details of the model comparison process.

### A.1. Process and Common Parameters

Common to all the models, the audio-only and visual-only trials were modelled with a mean, slope, and noise parameter (6 parameters total). Responses were modelled as draws from a truncated normal distribution peaked around the stimulus center times the slope plus the mean (like a simple linear regression between stimulus and response). For the audio-only trials, this truncated normal distribution had a standard deviation equal to the noise parameter. For the visual-only trials, the standard deviation was the noise parameter times the standard deviation of the displayed visual stimulus. The posterior vector of these six parameters were sampled by slice sampling.

For models with additional parameters, a draw was taken from that model’s prior for each sample. For each model, the sampled parameters (posterior on the unimodal data and any prior draws for the specific model) were used to calculate the joint probability of the audio-visual data. This joint probability was averaged over all samples, within each model, to get a score for the model and select the one that was the best in the Bayesian sense.

### A.2. Unique Parameters, Priors, and Computations

#### A.2.1. Optimal combination

Precision for each cue on each trial was calculated as each cue’s standard deviation to the power of negative 2. For this calculation, the visual cue’s actual standard deviation was used (not the one from fitting the visual-only data) since it intended to represent true optimal performance. Weights were taken as the relative reliability (each cue’s precision divided by the total precision) of the two cues. Since these precisions and weights were deterministic, there were no additional parameters to sample. The response was modelled as a truncated normal distribution with its peak at the sum of each single-cue aspect’s modal response (the stimulus times the slope plus the mean) times its weight. The variance of this truncated normal was the variance of the audio cue times its weight.

#### A.2.2. Biased weighting

This model is the same as above except the visual-only precision was multiplied by a draw from a Gamma(2,1) distribution.

#### A.2.3. Single weighting

This model is the same as the optimal model except the audio weight was a random draw from a Beta(2,2) distribution and the visual weight was one minus the audio weight. On each sample, the same weight was used for all trials (c.f. above where different visual-only aspects of the audio-visual stimulu, with their varying noise levels, led to different weights for every trial).

#### A.2.4. Switching

A switching weight was randomly sampled as Uniform(0,1). We then calculated the probability of the response as if it were an audio-only trial, then again as if it were a visual-only trial. The probability of the response was the as-if-visual probability times the switching weight plus the as-if-audio probability times one minus the switching weight.

#### A.2.5. Visual only

This model had no additional parameters. Probabilities were calculated as if they were additional visual-only trials (truncated normal with a peak at the slope times the stimulus plus the mean, with the true standard deviation times the noise parameter).

#### A.2.6. Audio only

This model also had no additional parameters. Probabilities were calculated as if they were additional audio-only trials (truncated normal with a peak at the slope times the stimulus plus the mean, with the noise parameter as a standard deviation).

